## Supplementary Information for "Fronto-motor circuits linked to effort-based decision-making and apathy in healthy subjects"

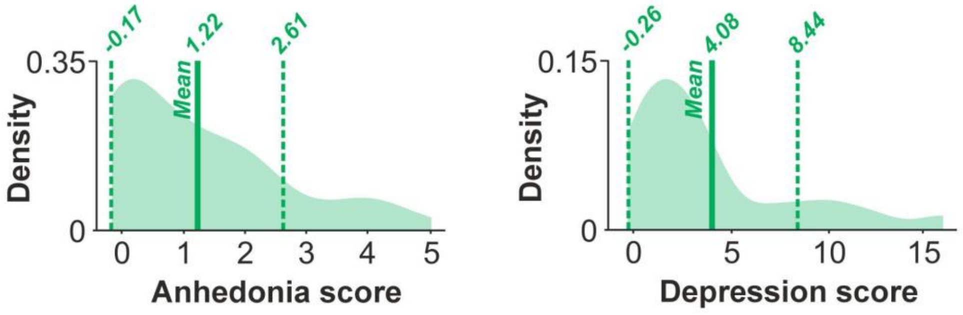


**Supplementary Figure 1: Distribution of anhedonia and depression scores.** We conducted a comprehensive neuropsychological assessment to quantify apathy scores, anhedonia scores and depression scores in our cohort of 45 healthy subjects. Apathy scores were assessed using the extended version of the Lille Apathy Rating Scale (LARS-e)[1], while anhedonia and depression were measured using the Snaith-Hamilton Pleasure Scale (SHAPS) and the Depression Anxiety Stress Scales (DASS), respectively. The figure presents density distributions of anhedonia and depression scores within the cohort. Anhedonia scores ranged from 0 to 5, with a mean of 1.22. Depression scores ranged from 0 to 16, with a mean of 4.08. These data reveal a broad range of anhedonia and depression levels across subjects, underscoring the importance of controlling for these variables through partial correlation analyses, as they may covary with apathy levels.

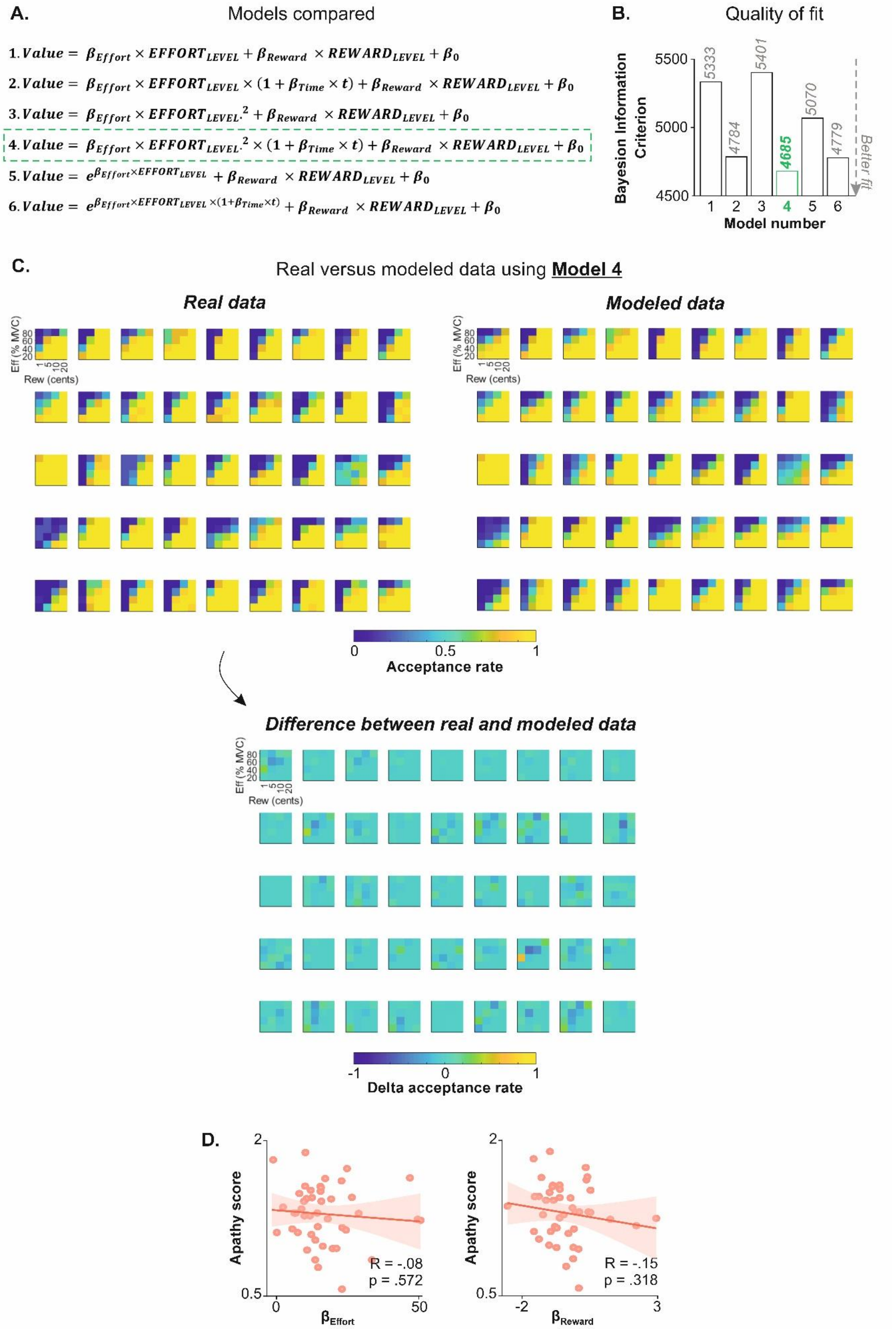


**Supplementary Figure 2: Computational modeling of decision behavior. A. Model comparison.** This panel shows the models tested in our study, following the same procedure as LeHeron et al., 2018[2]. Based on prior research[2–4], we evaluated candidate models of value computation using standard methods – minimization of the Bayesian Information Criterion (BIC, see B) and visual inspection of model fits (see C and Figure 1.B). The models differed in cost functions and whether they included a $\beta_{Time}$ parameter, which linearly modulated the cost term over trial number[4]. Specifically, Model 1 used a linear cost function without the $\beta_{Time}$ parameter, Model 2 used a linear cost function with $\beta_{Time}$, Model 3 used a quadratic cost function without $\beta_{Time}$, Model 4 used a quadratic cost function with $\beta_{Time}$, Model 5 used an exponential cost function without $\beta_{Time}$, and Model 6 used an exponential cost function with $\beta_{Time}$. **B. Model fit quality.** This panel presents the BIC for each model. Model 4, featuring a quadratic cost function and a $\beta_{Time}$ parameter, provided the best fit with the lowest BIC. This result aligns with previous studies[2,4], which also showed that quadratic cost functions with a $\beta_{Time}$ parameter gave the best fit for this type of task. **C. Comparison of real and modeled data.** The top left maps show the actual acceptance rates (color-coded from low (blue) to high (yellow) acceptance rates) in the effort-based decision-making task for each of the 45 subjects. The x-axis represents reward levels (1–20 cents), and the y-axis represents effort levels (20–80% MVC). The top right maps show modeled data using Model 4 for each subject. The bottom maps display the differences between real and modeled data, demonstrating that Model 4 closely fits subject-level data. These results complement the average data fit shown in Figure 1.B. **D. Apathy scores and computational parameters of effort and reward valuation capture different dimensions of behaviour.** While the computational parameters offer an objective, model-based way to capture variability in effort-based decision-making processes, apathy scores were not significantly correlated with $\beta_{Effort}$ and $\beta_{Reward}$ parameters. A Bayes Factor (BF) analysis showed that the $BF_{01}$ for these correlations were 2.97 and 2.24, respectively, indicating a higher likelihood of no correlation, consistent with prior work[4]. This indicates that effort-based decision-making offers a complementary approach that evaluates distinct dimensions of motivated behavior, not captured by apathy questionnaires.

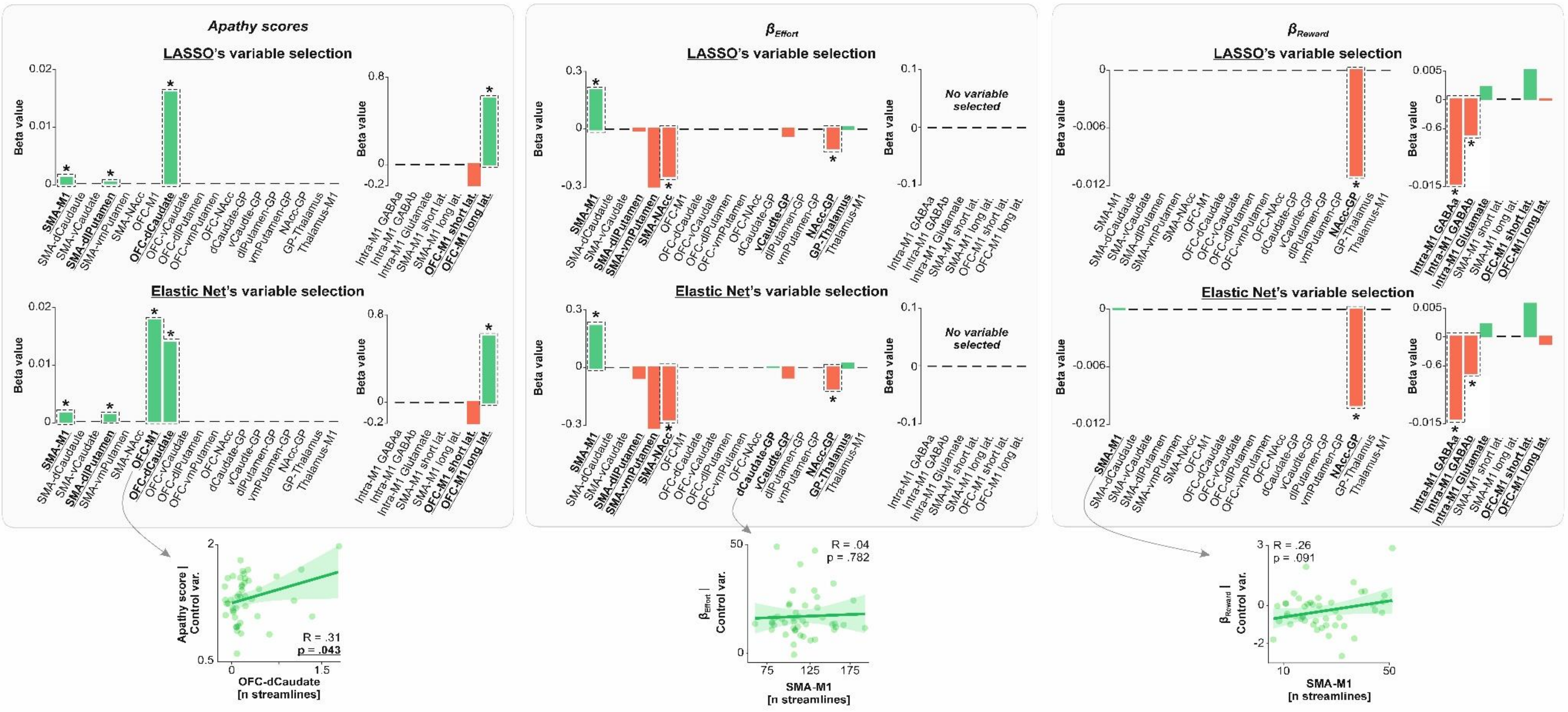


**Supplementary Figure 3: An Elastic Net regression reproduced the same results as the LASSO regression**. This figure presents the results of the Elastic Net regression analysis, which was conducted to address the potential risk of false negatives that may occur when using the conservative LASSO regression. As described in the main text, LASSO is a penalized least squares method that reduces the risk of false positives by selecting only relevant independent variables, excluding non-relevant variables by assigning them a regression coefficient of zero. While highly effective, LASSO's conservative nature may overlook relevant variables. To address this, we performed a less conservative Elastic Net regression. The results of the Elastic Net regression reproduced the main findings of the LASSO analysis, confirming the robustness of our results. Additionally, a complementary Bayes Factor Analysis provided further evidence for an absence of correlations between the apathy scores and the independent variables not selected by LASSO or Elastic Net regressions. Overall, across all analyses, LASSO regressions did not select 52 independent variables, providing beta coefficients equal to 0. The Elastic Net regression yielded highly similar results, excluding 49 independent variables. For $\beta_{Effort}$ and $\beta_{Reward}$, the Elastic Net regression selected two additional variables (*i.e.*, the number of streamlines in dCaudate-GP and SMA-M1 tracts, respectively), but neither achieved statistical significance when assessed via partial correlation analysis (bottom panels of the figure; p = .782 and .091, respectively). For apathy scores, the Elastic Net regression identified the OFC-M1 tract as an additional independent variable covarying with apathy. Partial correlation analysis showed a statistically significant association between apathy scores and OFC-M1 structural connectivity (p = .043). However, this finding should be interpreted with caution: the correlation is weak, is based on limited variance due to the low anatomical density of projections from OFC to M1 (see x-axis values), and lacks support from effective connectivity data obtained using TMS, where no significant OFC-M1 correlation was found (see Supplementary Figure 4). Collectively, these results highlight the robustness of the primary findings, particularly regarding potential false negatives when using LASSO regression.

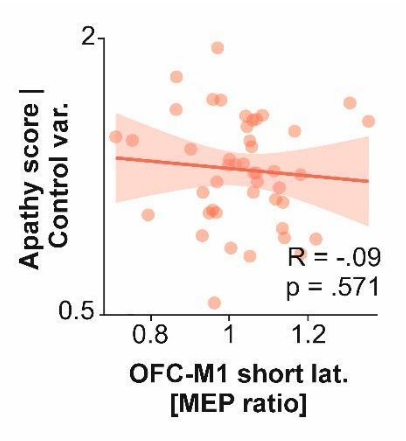


**Supplementary Figure 4: Apathy scores are not significantly correlated with effective connectivity in the short-latency OFC-M1 circuit.** The LASSO regression revealed two circuits (short- and long-latency OFC-M1) with non-zero coefficients (LASSO's β coefficients = -0.23 and 0.61, respectively; Figure 3). However, the partial correlation analysis showed that apathy scores were not significantly correlated with effective connectivity in the short-latency OFC-M1 circuit (R = -0.09, p = .571). The partial correlation only confirmed a significant positive correlation between apathy scores and the MEP ratio for the long-latency OFC-M1 circuit variable (R = 0.38, p = .014; Figure 3), indicating that higher apathy scores are associated with a stronger facilitatory influence of OFC on M1 specifically through this long-latency circuit.

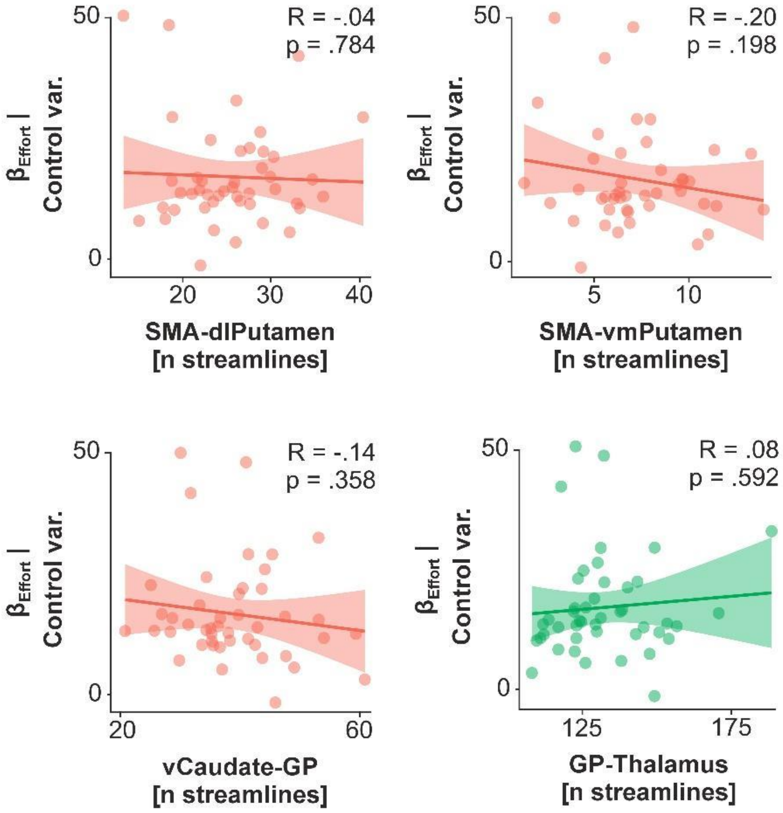


**Supplementary Figure 5: Effort valuation is not significantly correlated with structural connectivity in SMA-dlPutamen, SMA-vmPutamen, vCaudate-GP and GP-Thalamus circuits.** The LASSO regression identified seven tracts with non-zero coefficients: SMA-M1 (LASSO's β = 0.21), SMA-dlPutamen (β = -0.02), SMA-vmPutamen (β = -0.31), SMA-NAcc (β = -0.25), vCaudate-GP (β = -0.04), NAcc-GP (β = -0.11) and GP-Thalamus (β = 0.02; Figure 4). However, as evident in this figure, the partial correlation analysis showed that $\beta_{Effort}$ was not significantly correlated with structural connectivity in SMA-dlPutamen, SMA-vmPutamen, vCaudate-GP and GP-Thalamus circuits (p-values range = [.198 .784]). Partial correlation analyses showed that correlations between $\beta_{Effort}$ and the number of streamlines was only significant for three of these seven tracts, namely SMA-M1 (R = 0.42, p = .006), SMA-NAcc (R = -0.32, p = .042) and NAcc-GP circuits (R = -0.41, p = .007; Figure 4 in the main document).

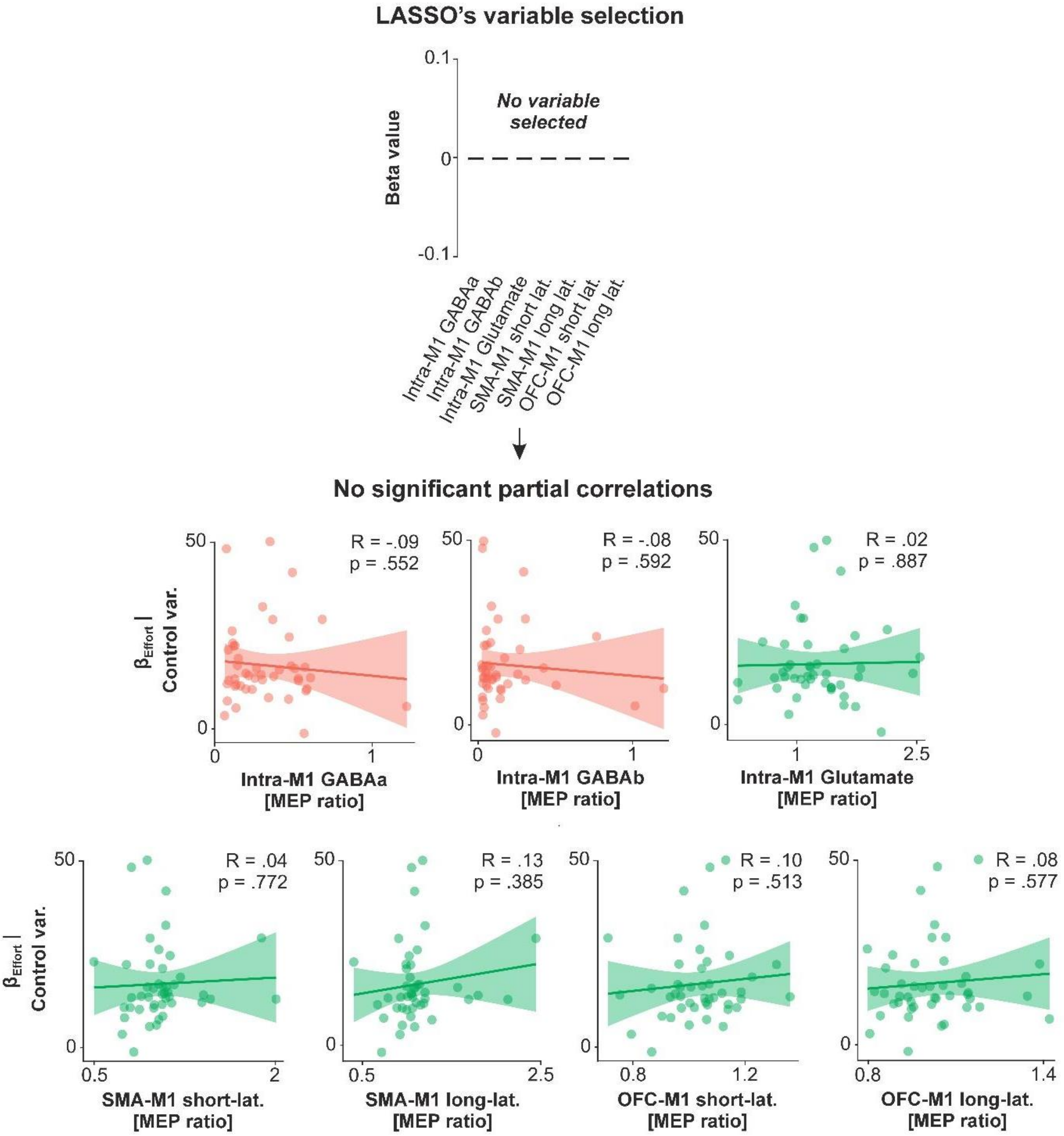


**Supplementary Figure 6: Effort valuation was not significantly correlated with effective connectivity in the investigated circuits.** We applied LASSO regression using $\beta_{Effort}$ as the dependent variable and all TMS effective connectivity data (*i.e.*, all MEP ratios) as independent variables. The regression yielded zero coefficients for all circuits (*i.e.*, all $\beta = 0$), signifying a lack of association between $\beta_{Effort}$ and effective connectivity in the investigated circuits (top graph). Partial correlations confirmed the absence of significant partial correlation between $\beta_{Effort}$ and all effective connectivity data (p-values range = [.385 .887]).

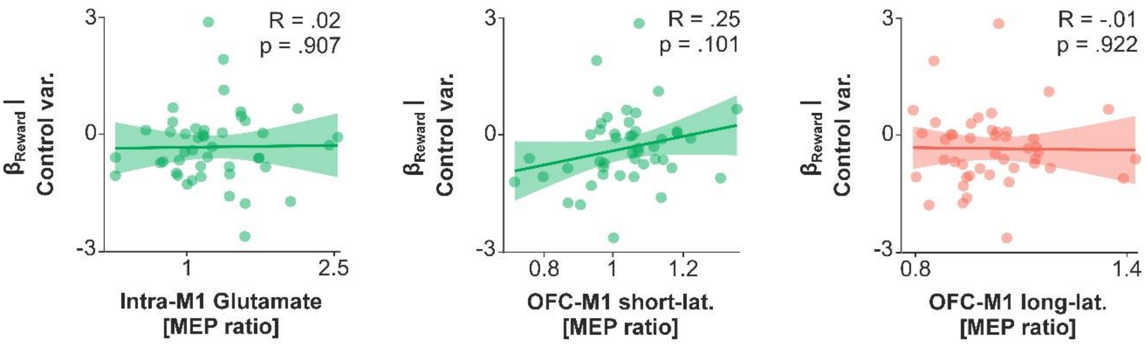


**Supplementary Figure 7: Reward valuation is not significantly correlated with effective connectivity in intra-M1 glutamatergic, and OFC-M1 short- and long-latency circuits.** The LASSO regression identified five circuits with non-zero coefficients (Figure 6): intra-M1 GABAa (LASSO's β = -0.015), intra-M1 GABAb (β = -0.006), intra-M1 glutamatergic (β = 0.002), short-latency OFC-M1 (β = 0.005) and long-latency OFC-M1 circuits (β = -0.0004). However, as evident in this figure, the partial correlation analysis showed that $\beta_{Reward}$ was not significantly correlated with effective connectivity in intra-M1 glutamatergic, and OFC-M1 short- and long-latency circuits (p-values range = [.101 .922]). Partial correlation analyses showed that correlations between $\beta_{Effort}$ and MEP ratios were only significant for two of these circuits, namely intra-M1 GABAa (R = -0.50, p = .0007) and intra-M1 GABAb circuits (R = -0.31, p = .047; Figure 6).

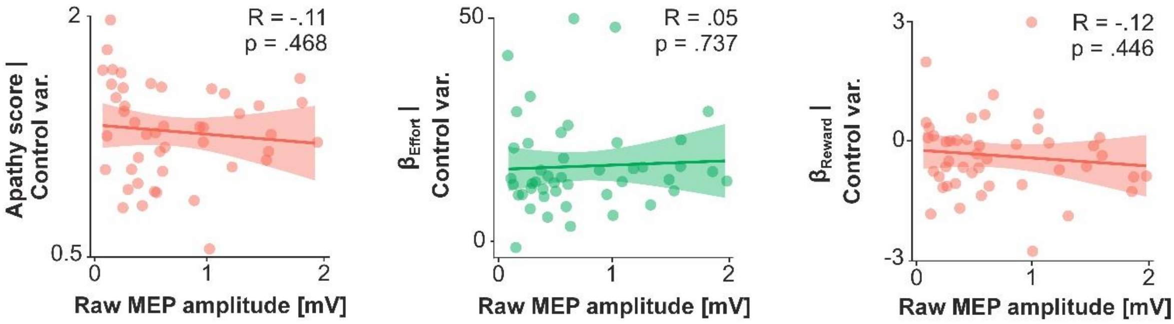


**Supplementary Figure 8: Apathy scores, effort valuation and reward valuation are not linked to M1 net output.** We performed partial correlations with apathy scores, $\beta_{Effort}$, and $\beta_{Reward}$ as dependent variables and test MEP amplitudes (obtained with a single test stimulation over M1), a proxy for M1 net output[5,6], as the independent variable. The analysis showed no significant correlation between apathy scores (R = -0.11, p = .468), $\beta_{Effort}$ (R = 0.05, p = .737), $\beta_{Reward}$ (R = -0.12, p = .446) and test MEP amplitudes. Bayes Factor computation indicated a higher likelihood of no correlation ($BF_{01}$ averaged 2.59 ± 0.34, range = [2.12 - 3.25]). Thus, despite correlations between apathy scores, effort valuation, reward valuation, and connectivity in circuits projecting to M1, these components of apathy are not linked to M1 net output alone. This analysis is also an important methodological control as it shows that the relationships between apathy scores, $\beta_{Effort}$, $\beta_{Reward}$, and MEP ratios are not due to associations with the test MEP amplitudes exploited to compute these ratios.

### Supplementary information references
